# Growing a cystic fibrosis-relevant polymicrobial biofilm to probe community phenotypes

**DOI:** 10.1101/2024.01.26.577445

**Authors:** Sarah Poirier, Fabrice Jean-Pierre

## Abstract

Most *in vitro* models lack the capacity to fully probe bacterial phenotypes emerging from the complex interactions observed in real-life environments. This is particularly true in the context of hard-to-treat chronic and polymicrobial biofilm-based infections detected in the airways of persons with cystic fibrosis (pwCF), a multiorgan genetic disease. While multiple microbiome studies have defined the microbial compositions detected in the airway of pwCF, no *in vitro* models thus far had fully integrated critical cystic fibrosis (CF)-relevant lung features. Therefore, a significant knowledge-gap in our capacity to investigate the mechanisms driving the pathogenesis of mixed species CF lung infections remained. To tackle this challenge, we have built a four-species microbial community model including *Pseudomonas aeruginosa, Staphylococcus aureus, Streptococcus sanguinis*, and *Prevotella melaninogenica* grown in CF-like conditions. Through the utilization of this system, clinically relevant phenotypes such as antimicrobial (Abx) recalcitrance of several pathogens were observed and explored at the molecular level. The usefulness of this *in vitro* model resides in its standardized workflow that can facilitate the study of interspecies interactions in the context of chronic CF lung infections.

**SUMMARY:** In this protocol, we describe a cystic fibrosis (CF)-lung relevant four-species polymicrobial biofilm model that can be used to explore the impact of bacterial interspecies interactions.

## INTRODUCTION

Strategies aimed at eradicating disease-causing microbes such as the ones detected in the cystic fibrosis (CF) airway, a multiorgan genetic disease, often fail.^1^ That is, the presence of resilient biofilm-like microbial communities growing in the mucus-rich CF lung environment can cause chronic infections spanning over multiple decades.^2^ Furthermore, although the utilization of front-line antimicrobials (Abx) and modulators has resulted in improved outcomes in persons with CF (pwCF), the “one bug, one infection” clinical approach− typically employed against canonical CF pathogens such as *Pseudomonas aeruginosa* and *Staphylococcus aureus*− has been ineffective in resolving hard-to-treat infections detected in the lungs of these individuals.^3-7^

Over the last two decades, multiple studies have changed our understanding of chronic CF lung disease. That is, reports indicate that infections detected in the airways of pwCF are not solely caused by a single pathogen but are rather polymicrobial in nature.^8^ Furthermore, although the pathogenesis of mixed species CF lung infections is still poorly understood, clinical evidence shows that pwCF that are co-infected with both *S. aureus* and *P. aeruginosa* in their airways have worsened lung function than individuals colonized by either of these two pathogens.^9^

Interspecies interactions among CF pathogens are hypothesized to be a reason as to why polymicrobial biofilm-based infections are not readily eradicated through the utilization of CF therapeutics, ultimately impacting patient outcomes.^3^ Supporting this, *in vitro* studies have demonstrated that interactions between microbes such as *P. aeruginosa, S. aureus* and others can impact clinically relevant phenotypes such as Abx responsiveness to front-line CF drugs, including vancomycin and tobramycin.^6,10,11^ Therefore, revisiting the current treatment strategies aimed at eradicating CF pathogens in the context of mixed species infections remain to be achieved.

The gold standard in the management of microbial-based CF lung infections heavily relies on the utilization of antimicrobial susceptibility testing (AST) to guide clinical interventions.^12^ However, AST is typically done using bacterial monocultures grown in rich and well-mixed cultures, which is not reflective of the microbial growth conditions detected in the CF airway.^3,13^ Given the significant disconnect between patient outcomes and treatment success, clinical reports now advocate for the development of novel approaches integrating critical features of the CF lung.^12,14^

While remaining informative, several studies have attempted to develop CF-relevant *in vitro* systems. However, these models (i) do not entirely reflect the polymicrobial and biofilm-like nature of the CF lung and, (ii) do not use nutritional conditions approximating the ones detected in the CF airway.^15,16^ Through mining of large CF-lung derived 16S rRNA gene data sets and computation, Jean-Pierre and colleagues recently developed an *in vitro* co-culture model integrating all the above-mentioned features.^10^ This system includes *P. aeruginosa, S. aureus, Streptococcus* spp. and *Prevotella* spp. grown in CF airway-like conditions and stable for up to 14 days. They also reported community-specific growth of *Prevotella* spp. and several changes in Abx responsiveness these CF pathogens.^10^ This tractable *in vitro* system also offered the possibility to delve into mechanistically focused questions related to Abx recalcitrance of a variant of *P. aeruginosa* frequently detected in the airways of pwCF.^10^ Therefore, the goal of this protocol is to provide the CF research community with a detailed protocol of this *in vitro* co-culture system, to probe polymicrobial-specific phenotypes.

## PROTOCOL

### A. Preparing Artificial Sputum Medium

<u>Note</u>: Throughout this protocol, artificial sputum medium (ASM) is defined as the CF-relevant medium as described previously by Turner and colleagues,^17^ which includes the presence of mucin.

Below, we describe the steps to prepare this medium. See **Table of Materials** to prepare stock solutions and storage conditions. This version of ASM differs from Turner *et al*.,^7^ as the buffering power in the original version does not allow for a stable pH over time. That is, fermentation activity carried out by *Streptococcus* spp. and anaerobes like *Prevotella* spp. results in the acidification of the growth medium (unpublished observations). Therefore, the MOPS concentration has been changed from 10 mM to 100 mM.

#### 1. 2X ASM base stock preparation

<u>Note</u>: Preparing a 2X ASM base (ASMb) stock solution facilitates the addition of components such as mucin, agar, water, etc. without impacting the final concentration of molecules present in ASM.^10,18^ The 2X ASMb can be prepared in advance and stored at 4°C for up to two weeks, in a dark place. If the medium turns yellow, discard.

In a clean beaker containing 400 mL of diH_2_O, add:

- 6.50 mL NaH_2_PO_4_*H_2_O stock
- 6.25 mL Na_2_HPO_4_ stock
- 348 μL KNO_3_ stock
- 1.084 mL K_2_SO_4_ stock
- 4.0 g Yeast Synthetic dropout without Tryptophan
- 3.032 g NaCl
- 20.92 g MOPS
- 1.116 g KCl
- 0.124 g NH_4_Cl
- 9.30 mL L-Lactic acid stock
- 2.70 mL glucose stock
- 1.75 mL CaCl_2_*2H_2_O stock
- 1.20 mL *N*-acetylglucosamine
- mL FeSO_4_*7H_2_O
- 660 μL Tryptophan stock
- 606 μL MgCl_2_*6H_2_O
- 0.60 g deoxyribonucleic acid frosm herring sperm
- 218 μL 1,2-dioleoyl-*sn*-glycero-3-phosphocholine (DOPC)

Once all the components have been added, adjust the pH to 6.80 with the 5M NaOH stock. At the end, complete to 500 mL with diH_2_O, filter sterilize on a 0.22 μm, and store at 4°C.

#### 2. Mucin stock preparation

To prepare a 10 mg/mL (2x) mucin stock, using a clean glass bottle containing a magnetic stir bar and 250 mL of diH_2_O, add:

- 5.0 g of mucin from porcine stomach – Type II
- Mix the suspension for 10 minutes.
- Complete to 500 mL with diH_2_O.
- Sterilize by autoclaving at 121°C, for 15 minutes.
- Keep at 4°C until utilization.

#### 3. Reconstituting the ASM

The day of the experiment:

- Mix a 1:1 ratio of the 2X ASMb stock and the 10 mg/mL mucin stock. This will be ASM, your working medium stock.
- Vortex thoroughly before use.

### B. Preparing polymicrobial community selective media

Below are the quantities necessary to prepare a volume of 1L of media. See **Table of Materials** for details.

- Mannitol Salt Agar (MSA) and *Pseudomonas* Isolation Agar (PIA).
  - Follow manufacturer’s recommendations.
- *Prevotella* Selective Agar (PSA)
  - 30.0 g tryptic soy broth (TSB)
  - 15.0 g agar
  - 5.00 g Yeast extract
  - 0.50 g L-Cysteine hydrochloride
  - 10.0 mL hemin stock
  - 100 μL menadione stock
  - Add 940 mL diH_2_O
  - Sterilize by autoclaving at 121°C for 15 minutes.
  - When cooled down (i.e., you can hold the bottle with your bare hands), add:
    - 50.0 mL defribrinated sheep’s blood
    - 2.00 mL of kanamycin stock (final concentration is 100 μg/mL)
    - 150 μL of vancomycin stock (final concentration is 7.5 μg/mL)
    - 500 μL of polymixin B stock (final concentration is 5 μg/mL)
- *Streptococcus* Selective Agar (SSA)
  - 30.0 g TSB
  - 15.0 g agar
  - Add 950 mL diH_2_O
  - Sterilize by autoclaving at 121°C for 15 minutes.
  - When cooled down (i.e., you can hold the bottle with your bare hands), add:
    - 50.0 mL defribrinated sheep’s blood
    - mL polymixin B stock (final concentration is 10 μg/mL)
    - mL oxolinic acid stock (final concentration is 10 μg/mL)
- Pour in plates and keep at 4°C for a maximum of 1 month. Before utilisation, let the plates dry up at room temperature up to 24 hr in advance before they are inoculated.

<u>Note</u>: PSA has been validated with *P. melaninogenica* and *Prevotella intermedia*. SSA has been validated with *S. sanguinis*, and the *Streptococcus* milleri group (*Streptococcus constellatus, Streptococcus intermedius* and *Streptococcus anginosus*, denoted as SMG). It is strongly recommended to *de novo* validate these selective media if new strains are to be tested.

### C. Bacterial liquid media preparation and growth conditions

- To routinely grow *P. aeruginosa* and *S. aureus*, use a sterile rich medium (e.g., lysis broth (LB) or TSB).
  - Start liquid overnights by picking a single colony previously grown on an LB/TSB agar plate for 20-24 hr at 37°C.
  - Grow broth cultures at 37°C overnight with shaking at 250 rpm for 16-18 hr.
- To grow *Streptococcus* spp., use sterile Todd Hewitt Broth supplemented with 0.5% yeast extract (THB-YE).
  - Start liquid overnight from a single colony previously grown on a TSB agar plate supplemented with 5% defibrinated sheep’s blood (blood agar plate) for 24 hr at 37°C + 5% CO_2_.
  - Grow the THB-YE cultures anaerobically or at 37°C + 5% CO_2_ overnight <u>without shaking</u> for 18-22 hr.
- To grow *Prevotella* spp. (*Prevotella* growth medium – PGM), prepare 1L of medium by adding:
  - 30.0 g TSB
  - 5.00 g yeast extract
  - 0.50 g L-cysteine hydrochloride
  - 10.0 mL Hemin stock
  - 100 μl menadione stock
  - Add 990 mL diH_2_O
  - Sterilize by autoclaving at 121°C for 15 minutes.
  - Wrap the bottle in aluminum foil to protect from light.
  - Grow *Prevotella* spp. colonies on a blood agar plate incubated anaerobically at 37°C for 48 hr.
  - Inoculate a single colony in PGM and grow overnight <u>without shaking</u> at 37°C anaerobically for 24 hr.

<u>Note</u>: PGM can be kept at room temperature for up to two weeks. Performing sterility tests on each growth medium is <u>strongly recommended</u> before use. If not using an anaerobic chamber, it might be necessary to start liquid overnights of *Prevotella* spp. from a patch of colonies.

### D. Co-culture experiment preparation

<u>Note</u>: **Figure 1** depicts a summarized experimental workflow. Setting up the experiment can be done in oxic conditions if the preparation time takes less than 1 hr. If it is expected to take longer than that, it is strongly recommended to perform the experiment using an anaerobic chamber (with 10% CO_2_, 10% H_2_, and 80% N_2_ mixed gas). Anaerobic jars (e.g., AnaeroPack™ 2.5L Rectangular Jars used with GasPaks - see **Table of Materials** for details), can also be used to incubate the experimental samples.

**Figure 1.**
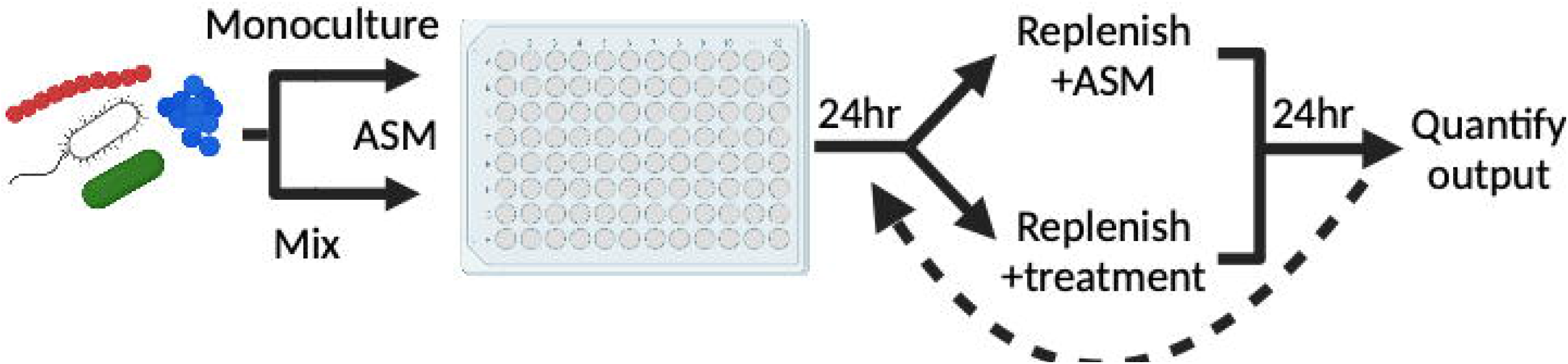
Experimental workflow. Growing monospecies and polymicrobial biofilms is done using a flat bottom 96-well plate into which microbes are cultivated in ASM anaerobically for 24 hr at 37°C. After the initial incubation step, non-attached cells are aspirated, and pre-formed biofilms are either replenished with fresh ASM or treated with molecules of interest. This step can be repeated on many iterations depending on the hypothesis to be tested. At the end of the experiment, cells are collected, serially diluted, and plated for viable counts. Figure created with BioRender.

#### 1. Preparation of the monocultures and co-cultures

<u>Note:</u> Do all the centrifugation steps at 10,000 x *g* for 2 min. For the co-cultures, mix the bacterial samples just before inoculating the 96-well plates. Typically, a volume of 1 mL for *P. aeruginosa* and *S. aureus* strains are collected from the overnights. For *Streptococcus* spp. and *Prevotella* spp., the overnight volumes may vary between 2-4 mL. These volumes can be adjusted depending on the number of conditions to be tested.

- Using the bacterial liquid overnights:
  - For *P. aeruginosa* and *S. aureus* strains, wash twice using sterile PBS.
  - For *Streptococcus* spp. and *Prevotella* spp., wash once with sterile PBS.
    - Discard supernatants carefully as the pellets can easily be lost.
  - After washes, resuspend each pellet in 1 mL of 1X water diluted ASMb.
  - Do a 1:10 dilution of each sample in sterile PBS and measure the OD_600_.
  - Adjust all the cultures to an OD_600_ of 0.2 in ASM.
  - Using the OD_600_ = 0.2 suspensions, dilute the bacterial samples to a <u>final</u> OD_600_ of 0.01 (for each microbe) for the monocultures and co-cultures.
  - Vortex thoroughly for 5 sec.

<u>Example</u>: To prepare a total of 1 mL of *P. aeruginosa* monoculture ASM suspension at OD_600_ = 0.01, add 50 μl of *P. aeruginosa* at an OD_600_ = 0.2 to a volume of 950 μl ASM. For the co-culture, take 50 μl of *P. aeruginosa, S. aureus, Streptococcus* spp. and *Prevotella* spp. at an OD_600_ = 0.2 and add to 800 μl ASM.

- Add 100 μl of the monoculture and co-culture suspensions in three separate wells of a sterile plastic flat bottom 96-well plate.
- Incubate the plate for 24 hr at 37°C in anoxic conditions.

<u>Note</u>: Other oxygen tensions (e.g., microxia, normoxia) can also be used here. However, the model has been developed and validated with anoxic conditions. Confirm the starting inocula by serially diluting the mono and co-culture bacterial suspensions and plating on PIA, MSA, SSA and PSA is strongly recommended. The targeted starting concentration for each bacterial species is 1 × 10^7^ CFU/mL (*P. aeruginosa*), 3.5 × 10^6^ CFU/mL (*S. aureus*), 1.2 × 10^6^ CFU/mL (*Streptococcus* spp.) and 4.6 × 10^6^ CFU/mL (*Prevotella* spp.). Bacterial inocula can also be modified as described by Jean-Pierre and colleagues.^10^

- After 24 hr incubation, remove the unattached (planktonic) cells by aspiration using a multichannel pipette.
- Replenish the pre-formed biofilms with 100 μl of fresh ASM or the desired treatment (e.g., antibiotics, metabolites, etc.).
- Incubate the plate for an additional 24 hr at 37°C in anoxic conditions.

### E. Collecting and plating the samples

- After the additional 24 hr of incubation, aspirate the non-attached (planktonic) cells with a multichannel pipette.
  - The planktonic fraction can be kept, serially diluted, and plated on selective media, or discarded.
- Gently wash the biofilms twice using 125 μl of sterile PBS and discard volumes.
- Add 50 μl of sterile PBS and detach the biofilms by gently scraping the cells from the plate using a 96-pin replicator (see **Table of Materials** for item description).
- Transfer the resuspended biofilm cells in row A of a new sterile 96-well plate.
  - The 96-well plate will contain sterile PBS in rows B to H.
- Perform a 10X serial dilution of the planktonic (if needed) and biofilm fractions.
- Plate 3-5 μl of each dilution sample onto PIA, MSA, SSA and PSA.
- Let the inoculation spots dry.
- For *P. aeruginosa* and *S. aureus*:
  - Incubate at 37°C for 18-20 hr.
- For *Streptococcus* spp.
  - Incubate at 37°C in anoxic conditions for 24 hr.
- For *Prevotella* spp.
  - Incubate at 37°C in anoxic conditions for 48 hr.
- After incubation, count the colonies (∼10-35 per spot) and calculate the colony forming units (CFU) per mL.
- Plot the data using a visualization tool.

## REPRESENTATIVE RESULT

As represented in **Figure 2**, several phenotypes were reported including (**i**) a reduction in the number of *P. aeruginosa* and *S. aureus* viable cell counts when grown a mixed planktonic communities compared to monoculture, (**ii**) an increase in polymicrobial growth of *S. sanguinis* cells and, (**iii**) mixed community growth of *P. melaninogenica* as previously reported by Jean-Pierre and colleagues.^10^

**Figure 2.**
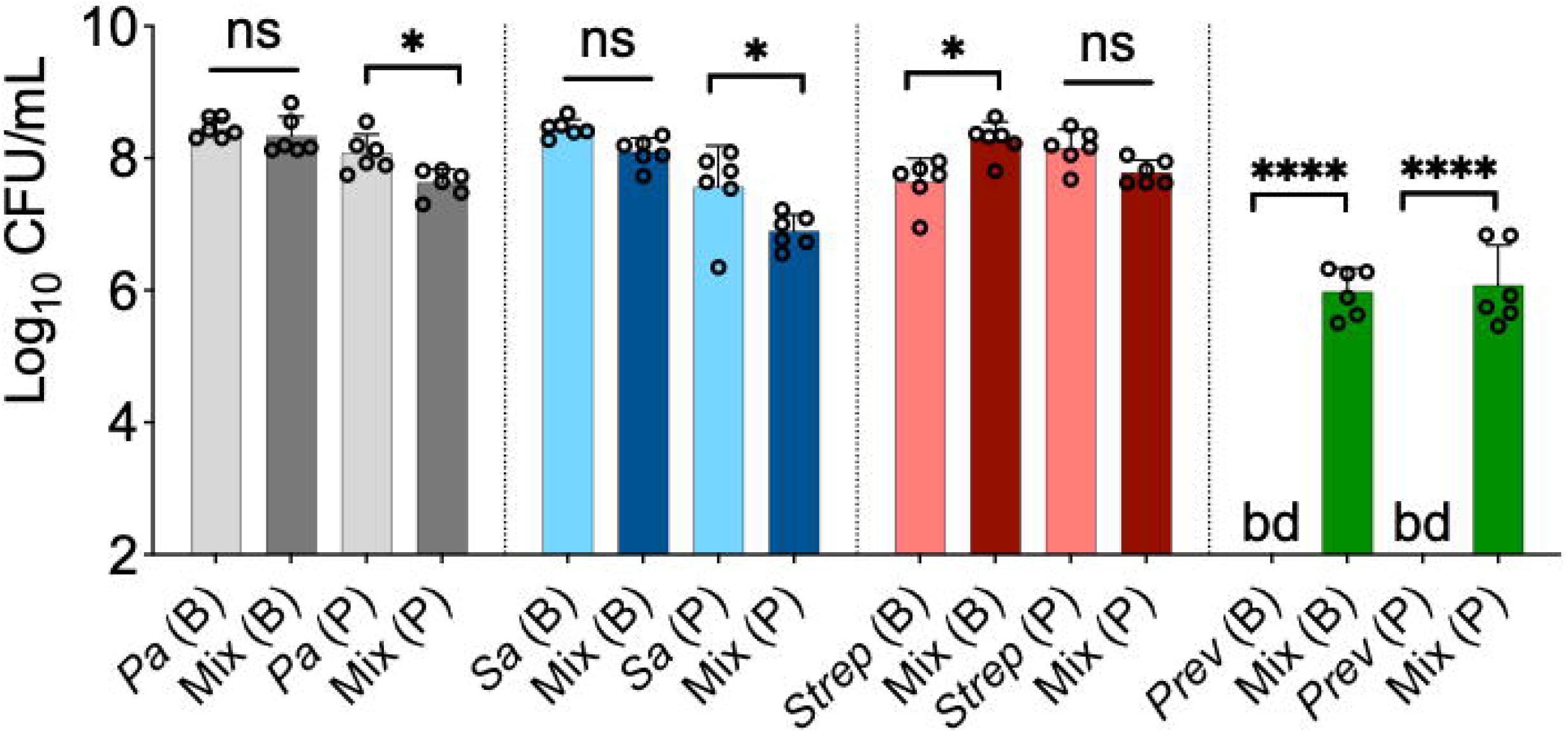
Detecting polymicrobial-specific phenotypes using *in vitro* modeling. Indicated are CFU/mL counts of *P. aeruginosa* (*Pa*), *S. aureus* (*Sa*), *S. sanguinis* (*Strep*) and, *P. melaninogenica* (*Prev*) bacterial species grown as a monoculture or in a mixed community (Mix) for biofilm (B) and planktonic (P) fractions. Represented data point presented in a column represents the average from at least three technical replicates performed at least on six different days (n = 6). Statistical analysis was performed using ordinary one-way analysis of variance (ANOVA) and Tukey’s multiple comparisons posttest with *, P < 0.05; ****, P < 0.0001, ns = non-significant. Error bars represent the standard deviation. bd = below detection. Figure modified from Jean-Pierre *et al*.,^10^ with permission (Creative Commons Attribution license).

Furthermore, as depicted in **Figure 1**, the *in vitro* system can also be used to test molecules of interest (e.g., Abx, metabolites, etc.) once the biofilm communities have been established. That is, microbes grown in co-culture have either shown (**i**) an increase in susceptibility (*P. aeruginosa* – **Figure 3A**), (**ii**) a decreased in sensitivity (*S. aureus* and *S. sanguinis* – **Figure 3B, 3C**) or (**iii**) no changes (*P. melaninogenica* – **Figure 3D**) in the presence of the CF drug tobramycin versus monoculture.

**Figure 3.**
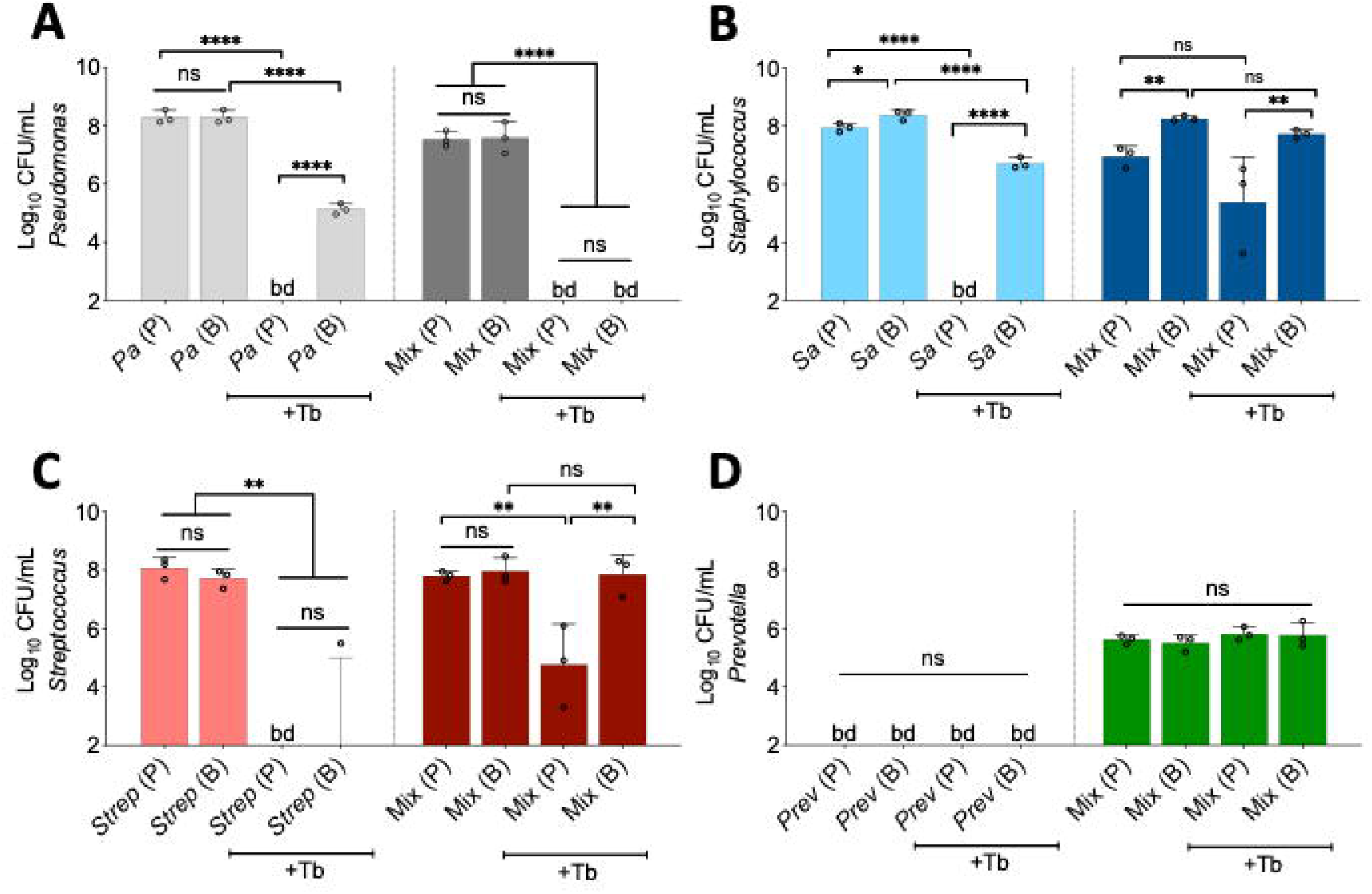
Interspecies interactions impact Abx susceptibility of CF pathogens. Indicated are CFU/mL counts of (**A**) *P. aeruginosa* (*Pa*), (**B**) *S. aureus* (*Sa*), (**C**) *S. sanguinis* (*Strep)*, and (**D**) *P. melaninogenica* (*Prev*) bacterial populations grown as a monoculture or in a polymicrobial (Mix) in the absence or presence of tobramycin (+Tb) at a concentration of 100 μg/mL. Represented data point presented in a column represents the average from at least three technical replicates performed at least on three different days (n = 3). Statistical analysis was performed using ordinary one-way analysis of variance (ANOVA) and Tukey’s multiple comparisons posttest with *, P < 0.05; **, P < 0.01, and ****, P < 0.0001. ns = non-significant. Error bars represent the standard deviation. bd = below detection. Figure modified from Jean-Pierre *et al*.,^10^ with permission (Creative Commons Attribution license).

## Table of Materials

Materials and stock solutions necessary to use the *in vitro* co-culture system.

## DISCUSSION

The usefulness of this clinically informed *in vitro* co-culture system resides in its capacity to detect polymicrobial-specific bacterial functions. That is, through the utilization of this model, we have reported novel microbial phenotypes ranging from community-specific growth of *Prevotella* spp. (**Figure 2**) to changes in susceptibility to front-line CF Abx of *P. aeruginosa, S. aureus* and *S. sanguinis* (**Figure 3**). Moreover, even though data presented in **Figures 2 & 3** are representative of laboratory strains, similar results have been obtained using various species (SMG and *P. intermedia*) and clinical isolates.

We argue that this co-culture system can be leveraged to test a large spectrum of CF-related questions ranging from virulence to microbe-microbe/host-microbe interactions. For example, Kesthely and colleagues have recently looked at changes in transcriptional responses of microbes grown in co-culture *versus* monocultures using the polymicrobial model described here.^19^ Furthermore, although the development of novel modulators has resulted in improved outcomes in pwCF, chronic microbial-based infections in their airways persist.^4,5,7^ Therefore, we propose that this *in vitro* community model represents an ideal platform to explore the mechanisms allowing these communities to thrive in the CF lung in the presence of these modulators.

The major technical hurdles encountered in using this model generally are (**i**) the preparation of the monocultures and co-cultures mixes and, (**ii**) cross-contamination between samples. For the former point, limiting the number of experimental conditions (i.e., identifying the minimal number of conditions necessary to test a hypothesis while including all appropriate controls in a rigorous and reproducible manner) is a strategy. Also, preparing the experiment in an anaerobic chamber can mitigate some of the issues that can arise while setting up the experiment in oxic conditions on the bench. For the latter point, including several controls as checkpoints is critical to alleviate the risks.

Finally, a limitation of this *in vitro* co-culture system is the absence of host-derived factors (e.g., immune cells, host-derived metabolites, etc.) that are known to be important in the context of CF lung disease.^20^ However, we argue that this polymicrobial model can be used as a starting point to identify mechanisms that can be further vetted in more *in vivo*-like systems.

## ACKNOWLEDGMENTS

We would like to acknowledge Dr. George A. O’Toole and Dr. Thomas H. Hampton for their significant role in the design and development of the *in vitro* polymicrobial community model. We thank Dr. Sophie Robitaille for helpful comments on the manuscript. This work was supported by a CFF grant JEAN21F0 to F.J-P.

